# Proteolytic cleavage of *Arabidopsis thaliana* phospho*enol*pyruvate carboxykinase-1 modifies its allosteric regulation

**DOI:** 10.1101/2020.07.29.226720

**Authors:** Bruno E. Rojas, Matías D. Hartman, Carlos M. Figueroa, Alberto A. Iglesias

**Affiliations:** Instituto de Agrobiotecnología del Litoral, UNL, CONICET, FBCB, Santa Fe, Argentina

**Keywords:** *Arabidopsis thaliana*, gluconeogenesis, metacaspase, phospho*enol*pyruvate carboxykinase, proteolysis, seedlings

## Abstract

Phospho*enol*pyruvate carboxykinase (PEPCK) plays a crucial role in gluconeogenesis. In this work, we analyze the proteolysis of *Arabidopsis thaliana* PEPCK1 (*Ath*PEPCK1) in germinating seedlings. We found that expression of *Ath*PEPCK1 peaks at 24-48 hours post-imbibition. Concomitantly, we observed shorter versions of *Ath*PEPCK1, putatively generated by metacaspase-9 (*Ath*MC9). To study the impact of *Ath*MC9 cleavage on the kinetic and regulatory properties of *Ath*PEPCK1, we produced truncated mutants based on the reported *Ath*MC9 cleavage sites. The Δ19 and Δ101 truncated mutants of *Ath*PEPCK1 showed similar kinetic parameters and the same quaternary structure than the WT. However, activation by malate and inhibition by glucose 6-phosphate were abolished in the Δ101 mutant. We propose that proteolysis of *Ath*PEPCK1 in germinating seedlings operates as a mechanism to adapt the sensitivity to allosteric regulation during the sink-to-source transition.

**Highlight:** This paper describes the effects of the N-terminal proteolytic cleavage on the kinetic and regulatory properties of *Arabidopsis thaliana* phospho*enol*pyruvate carboxykinase-1.

## Introduction

Phospho*enol*pyruvate (PEP) is a metabolic hub that connects various pathways, including glycolysis, gluconeogenesis, and metabolism of organic and amino acids (Chiba *et al*., 2015). Among the enzymes that metabolize PEP, PEPCK is particularly relevant, due to its numerous physiological functions (Leegood and Walker, 2003; Latorre-Muro *et al*., 2018; Wang and Dong, 2019). Based on the phosphate donor, PEPCKs are classified as ATP- (EC 4.1.1.49), GTP- (EC 4.1.1.32) or PPi-dependent (EC 4.1.1.38), all with different evolutionary origin (Matte *et al*., 1997; Fukuda *et al*., 2004; Aich and Delbaere, 2007; Chiba *et al*., 2015).

ATP-dependent PEPCK is found in bacteria, yeasts, and plants. This enzyme catalyzes the reversible decarboxylation of oxaloacetate (OAA) to form PEP, according to the reaction: OAA + ATP ⇆ PEP + ADP + CO_2_. Although this reaction is fully reversible *in vitro*, it is generally accepted that it proceeds towards OAA decarboxylation *in vivo* (Johnson *et al*., 2016). PEPCK requires two divalent cations to catalyze the reaction: one Mn^2+^ ion acts as an essential activating cofactor that promotes OAA decarboxylation and stabilizes the enolate ion during catalysis; while one Mg^2+^ cation forms the metal nucleotide complex that constitutes the active form of the substrate (Goldie and Sanwal, 1980; Burnell, 1986; Matte *et al*., 1997; Johnson *et al*., 2016).

ATP-dependent PEPCK has different physiological roles in plants: i) it is part of the CO_2_-concentrating mechanisms operating in C_4_ and CAM photosynthesis (Edwards *et al*., 1971; Reiskind and Bowes, 1991; Martín *et al*., 2011); ii) it participates in biotic and abiotic stress responses (Saez-Vasquez *et al*., 1995; Chen *et al*., 2000, 2002; Saito *et al*., 2008; Penfield *et al*., 2012; Choi *et al*., 2015); iii) it is involved in nitrogen and amino acid metabolism, especially during fruit development (Walker *et al*., 1999; Lea *et al*., 2001); and iv) it is involved in gluconeogenesis during seed germination, channeling carbon released from fatty acid reserves to form sugars, until the photosynthetic apparatus is fully developed (Rylott *et al*., 2003; Penfield *et al*., 2004; Malone *et al*., 2007; Graham, 2008; Eastmond *et al*., 2015).

The occurrence and regulatory effects of proteolysis on PEPCK have remained obscure. Walker *et al*. (1995) found that a discrete proteolytic cleavage at the N-terminus of PEPCK occurred in crude extracts from cucumber cotyledons and leaves of C_4_ and CAM species; however, proteolysis of the cucumber PEPCK did not significantly alter its activity (Walker and Leegood, 1995). Two PEPCK isoforms of different molecular mass (74 and 65 kDa) were found in *Ananas comosus* (pineapple) leaves. The shorter version was purified to homogeneity and biochemically characterized, but study of the large isoform was not possible because it was recalcitrant to purification (Martín *et al*., 2011). A large scale study conducted by Tsiatsiani *et al*. (2013) identified *Ath*PEPCK1 as a substrate of *Arabidopsis thaliana* (Arabidopsis) metacaspase-9 (*Ath*MC9). This cysteine protease cleaved *Ath*PEPCK1 at the N-terminus, which seemed to boost PEPCK activity. In crude extracts, PEPCK activity was reduced in the *mc9* mutant and increased in the MC9 overexpressing lines (Tsiatsiani *et al*., 2013).

Arabidopsis has two *ATP-PEPCK* genes, *pck1* (AT4G37870) and *pck2* (AT5G65690), encoding *Ath*PEPCK1 and *Ath*PEPCK2, respectively. We have recently reported the biochemical properties of these proteins, which are finely regulated by numerous metabolites. Mainly, they are inhibited by glucose 6-phosphate (Glc6P), shikimate, and inorganic pyrophosphate (PPi), and activated by malate (Rojas *et al*., 2019). In this work, we focused on the combined effects of allosteric regulation and proteolysis on the activity of *Ath*PEPCK1, which plays a critical role during seed germination (Rylott *et al*., 2003; Penfield *et al*., 2004). We used recombinant proteins to study in detail the biochemical effects of the *Ath*MC9-mediated cleavage of *Ath*PEPCK1. Our results show that proteolysis of the N-terminus modifies the allosteric regulation of *Ath*PEPCK1, which could have important metabolic implications during the sink-to-source transition associated with seedling development.

## Materials and methods

### Reagents

ATP, ADP, PEP, OAA, NADH, Glc6P, L-malic acid, pyruvate kinase, and L-malic dehydrogenase were from Sigma Aldrich. L-lactate dehydrogenase was from Roche. All other reagents were of the highest available quality.

### Plant material and growth conditions

All experiments were performed with Arabidopsis Col-0. The *pck1* mutant corresponds to the SALK_072899C T-DNA insertion. Seeds were disinfected with 70% (v/v) ethanol for 5 min, then treated with 10% (v/v) bleach for 10 min and washed thrice with sterile, distilled water.

Seeds were soaked in sterile 0.1% (w/v) agar and stratified at 4 °C in the dark for two days. In germinating assays, seeds were sown on a mesh soaked with 0.5 X Murashige and Skoog medium in 16 cm diameter Petri dishes and transferred to growth chambers (Raineri *et al*., 2016). In mature leaf assays, plants were grown on soil in 8 cm diameter x 7 cm height pots, one plant per pot. In all cases, plants were grown at 23 °C and 120 μmol m^−2^ s^−1^, with a long day photoperiod (16 h light and 8 h dark). The seeds employed were harvested, dried in darkness at room temperature, and stored at 4 °C. In all experiments, samples were taken, immediately frozen with liquid nitrogen, and stored at −80 °C until use.

### Protein extraction

Plant material was homogenized in a precooled mortar with liquid nitrogen. For denaturing protein extraction, 20 mg of fresh weight (FW) tissue was extracted with 200 μl of denaturing sample buffer, consisting of 2% (w/v) SDS, 20% (w/v) glycerol, 1.4 M 2-mercaptoethanol, 125 mM Tris-HCl pH 6.8 and 0.05% (w/v) Bromophenol Blue. After adding the buffer, samples were vortexed and heated for 5 min at 95 °C with agitation. Samples were cooled to room temperature and centrifuged at 21,000 x *g* for 10 min to separate the protein extract from tissue debris. For native protein extraction, 20 mg of FW tissue was extracted with 500 μl of native buffer, consisting in 100 mM Bicine-KOH pH 9.0, 10% (v/v) glycerol, 0.1% (v/v) Triton X-100, 5 mM 2-mercaptoethanol, 1 mM EDTA, 1 mM EGTA, 1 mM benzamidine, 1 mM ε-aminocapronic acid, 2 mM PMSF, 1X Set III protease cocktail (Merck, 539134), 1 mM NaF, 1 mM Na_2_MO_4_, and 1 mM Na_3_VO_4_. After adding the extraction buffer, samples were vortexed and incubated on ice for 10 min and then centrifuged at 21,000 x *g* for 10 min. Then, the protein extract was separated from tissue debris and transferred to a new tube.

### Production and purification of recombinant proteins

The production of recombinant *Ath*PEPCK1 was performed as previously described (Rojas *et al*., 2019). The sequence coding for *Ath*MC9 (At5g04200) was cloned using cDNA from Arabidopsis seedlings and inserted in frame with an N-terminal His_6_-tag between the *Bam*HI and *Eco*RI sites of the pETDuet-1 vector (Novagen). Protein expression was performed in *E. coli* BL21 (DE3) (Invitrogen) grown in LB medium supplemented with 100 μg ml^−1^ ampicillin and induced with 0.5 mM isopropyl-β-D-1-thiogalactopyranoside for 16 h at 18 °C with agitation. Cells were harvested by centrifugation and resuspended in lysis buffer [25 mM Tris-HCl pH 8.0, 300 mM NaCl, 5% (v/v) glycerol and 10 mM imidazole]. Cells were disrupted by sonication and centrifuged at 12,000 x *g* for 15 min at 4 °C. The crude extract was loaded on an IDA-Ni^2+^ column, previously equilibrated with lysis buffer. The recombinant protein was eluted with lysis buffer supplemented with 300 mM imidazole and 10% (v/v) glycerol.

### Production of antisera

Polyclonal antibodies against *Ath*PEPCK1 were raised in rabbits using the purified recombinant protein at the Centro de Medicina Comparada (ICIVET Litoral, CONICET-UNL, Argentina). To further increase the specificity, the antiserum was purified with *Ath*PEPCK1, following a previously described protocol (Fang, 2012). Polyclonal antibodies against *Titricum aestivum* NAD-dependent glyceraldehyde-3-phosphate dehydrogenase (*Tae*GAPDH) were produced by Piattoni *et al*. (2017).

### Protein methods

Total proteins were quantified with the Bradford assay (Bradford, 1976), using a standard curve constructed with bovine serum albumin.

Protein electrophoresis was performed under denaturing conditions (SDS-PAGE), according to the method described by Laemmli (1970). For immunodetection, proteins were transferred to 0.45 μm nitrocellulose membranes (Amersham) at 180 mA for 60 min. Membranes were incubated overnight at 4 °C with purified anti-*Ath*PEPCK1 antibodies diluted 1/1,000 and then incubated for 1 h at room temperature with goat anti-rabbit IgG H&L conjugated to horseradish peroxidase (Abcam, ab6721) diluted 1/10,000. Protein bands were revealed with SuperSignal West Pico Plus Chemiluminescent Substrate (Thermo Fischer Scientific), following the manufacturer’s instructions. Radiographic films (AGFA) were exposed between 2.5 and 5 min for the detection of *Ath*PEPCK1. Antibodies were stripped with 100 mM glycine pH 2.5 to analyze protein loading. Membranes were thoroughly washed and then incubated overnight with anti-*Tae*GAPDH antibodies diluted 1/5,000. All subsequent steps were performed as previously described. In this case, exposition times were between 30 s and 1 min.

### Enzyme activity assays

Recombinant *Ath*PEPCK1 and the Δ19 and Δ101 truncated mutants were assayed as previously described (Rojas *et al*., 2019). Carboxykinase activity of PEPCK in crude extracts was measured using 100 mM HEPES-NaOH pH 7.0, 4 mM 2-mercaptoethanol, 0.2 mM NADH, 4 mM MgCl_2_, 1 mM MnCl_2_, 100 mM KHCO_3_, 0.2 mM ADP, 10 mM PEP and 1 U malate dehydrogenase. Assays were performed in 250 μl at 30 °C and were corrected for phospho*enol*pyruvate carboxylase activity by omitting ADP from the reaction mixture, as previously done by Martin *et al*. (2007).

Recombinant *Ath*MC9 protein was assayed as described previously by Vercammen *et al*. (2004), with minor modifications. Reactions were done with 50 mM MES-KOH pH 5.5, 150 mM NaCl, 300 mM sucrose, 10 mM DTT, 0.3 μg μl^−1^ *Ath*MC9, and 0.15 μg μl^−1^ *Ath*PEPCK1. Aliquots were taken at different time intervals and analyzed by SDS-PAGE and PEPCK activity.

## Results

### Proteolysis of AthPEPCK1 in germinating seedlings

Transcriptomic data retrieved from the eFP browser (Winter *et al*., 2007) shows that the expression of *AthPEPCK1* and *AthPEPCK2* in germinating seedlings and mature leaves are considerably different (Fig. S1). *AthPEPCK1* transcripts are 620-fold higher than *AthPEPCK2* transcripts in germinating seedlings at 48 hours after imbibition (HAI; Fig. S1). *AthPEPCK1* transcripts are 10-fold higher in germinating seedlings at 48 HAI than in mature leaves, while *AthPEPCK2* transcripts are relatively low in germinating seedlings and below the detection limit in mature leaves (Fig. S1). Based on this information, our experiments were focused on *Ath*PEPCK1. To analyze the integrity of this protein at different developmental stages, we extracted proteins under denaturing conditions from germinating seedlings harvested at 48 HAI and mature leaves from 32-days-old rosettes (Fig. 1A). We found that *Ath*PEPCK1 is proteolyzed in germinating seedlings but not in mature leaves, although the amount of *Ath*PEPCK1 is lower in the latter (Fig. 1A). Some proteases can be active during protein extraction, even with sample buffer containing SDS (Plaxton, 2019). To discard the possibility that *Ath*PEPCK1 from germinating seedlings is degraded during the extraction, the sample buffer was supplemented with 2 M urea. Fig. S2 shows that proteolysis of *Ath*PEPCK1 occurs *in vivo* and is not an artifact of the extraction procedure.

**Fig. 1.**
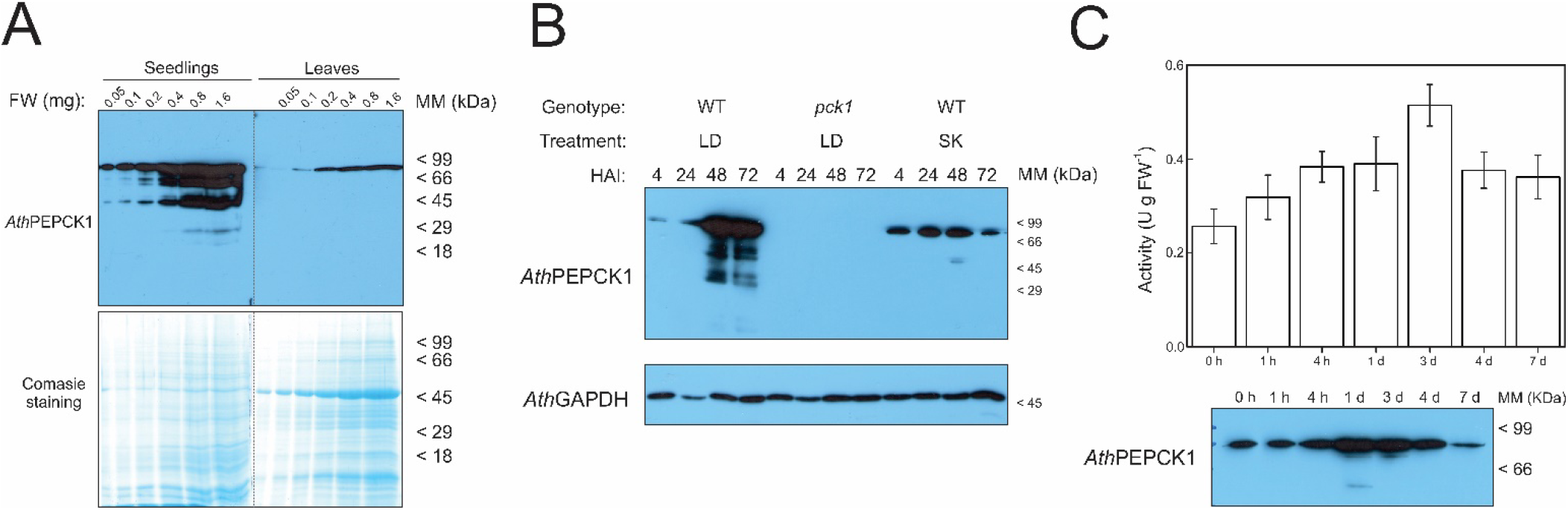
Proteolysis of *Ath*PEPCK1 in germinating seedlings. A) Analysis of *Ath*PEPCK1 in germinating seedlings and mature rosettes. Denatured protein extracts from Arabidopsis 48-HAI germinating seedlings (left) and 32-days-old rosettes (right) were resolved in a 12% SDS-PAGE (lower panel). Proteins were transferred to nitrocellulose membranes and immunodetected with anti-*Ath*PEPCK1 antiserum (upper panel), as described in the Materials and methods section. FW; amount of fresh weight tissue loaded in the gel. B) Time course of *Ath*PEPCK1 proteolysis during Arabidopsis germination. Western blot performed with anti-*Ath*PEPCK1 antiserum (upper panel) and load control with anti-*Tae*GAPDH antiserum (lower panel) of denatured protein extracts from Arabidopsis seedlings, obtained as described in the Materials and methods section. HAI; hours after imbibition. LD; long-day condition (16 h light and 8 h dark). SK; skotomorphogenesis (total darkness). C) PEPCK activity in Arabidopsis germinating seedlings. PEPCK carboxykinase activity was measured in crude extracts from Arabidopsis seedlings, as described in the Materials and methods section. Data are the mean ± standard error of four biological replicates.

Polyclonal antiserum raised against recombinant *Ath*PEPCK1 cross-reacts with *Ath*PEPCK2 (78.4% sequence identity, data not shown). Considering the differences in the relative abundances of *AthPEPCK1* and *AthPEPCK2* transcripts in germinating seedlings (Fig. S1), we assumed that the main isoform detected in Fig. 1A is *Ath*PEPCK1. To test our hypothesis, we analyzed protein extracts obtained under denaturing conditions from germinating seedlings of WT plants and the *pck1* knock-out mutant. Seedlings were grown under long-day conditions and samples were harvested at time intervals from 4 to 72 HAI. Fig. 1B shows that *Ath*PEPCK1 proteolysis peaked at 24-48 HAI in WT plants, whereas no protein bands were detected in the *pck1* mutant at any time point. The proteolysis of *Ath*PEPCK1 was also observed in WT seedlings grown in total dark (skotomorphogenesis), even though the amount of *Ath*PEPCK1 was significantly lower than in seedlings grown under long-day conditions (Fig. 1B).

Based on our results (Figs 1A, 1B, and S2) and the reports of PEPCK degradation in cucumber cotyledons (Walker *et al*., 1995), we optimized a protocol for the extraction of native proteins to assess PEPCK activity in germinating seedlings. Extraction was performed with a buffer at pH 9.0 (which diminished PEPCK proteolysis in protein extracts from cucumber; Walker & Leegood, 1995) supplemented with protease inhibitors, to avoid further degradation during the assays. We found that *Ath*PEPCK1 remained stable for at least 4 h after the extraction (Fig. S3) and PEPCK activity in crude extracts from Arabidopsis seedlings was in the range described by Malone *et al*. (2007). As shown in Fig. 1C, *Ath*PEPCK1 activity was highest at 4 days after imbibition (DAI), while the peak of protein expression was observed at 1 DAI. The maximum level of *Ath*PEPCK1 protein coincided with the onset of proteolysis, around 24-72 HAI (Figs 1B and 1C).

### Kinetic and regulatory effects on AthPEPCK1 triggered by AthMC9 cleavage

The multiple *Ath*PEPCK1 bands observed in crude extracts suggested partial proteolysis by a protease (Figs 1A-C and S2-S3). It has been previously reported that *Ath*PEPCK1 is cleaved at the N-terminus by *Ath*MC9 (Tsiatsiani *et al*., 2013). To test if *Ath*PEPCK1 is cleaved by *Ath*MC9, we incubated recombinant *Ath*PEPCK1 for 60 min with crude extracts from germinating seedlings harvested at 48 HAI at different pH values, as *Ath*MC9 is inactive at alkaline pH (Vercammen *et al*., 2004). We found that the cleavage of *Ath*PEPCK1 occurred at pH 5.5 and 7.0, whereas it was prevented at pH 9.0 (Fig. 2A).

**Fig. 2.**
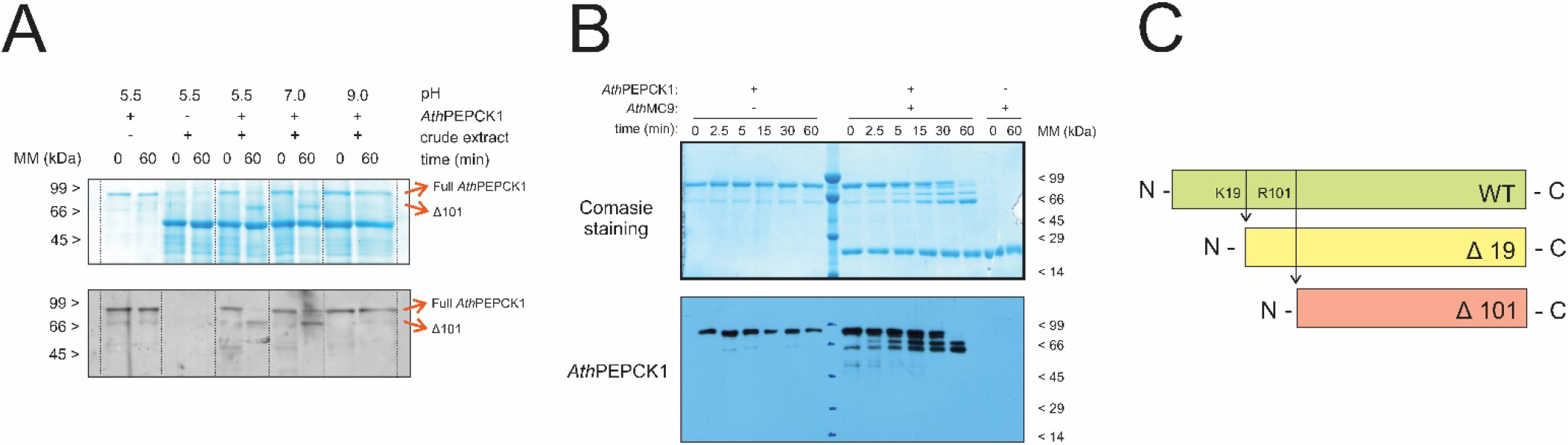
Analysis of *Ath*PEPCK1 cleavage by *Ath*MC9. A) Cleavage of *Ath*PEPCK1 by crude extracts from Arabidopsis seedlings. Recombinant *Ath*PEPCK1 was incubated with protein extracts from Arabidopsis seedlings in MES-NaOH pH 5.5, HEPES-NaOH pH 7.0, or Tricine-NaOH pH 9.0 (150 mM in all cases). Reactions were terminated by adding denaturing sample buffer, resolved on a 10% SDS-PAGE (upper panel), and immunodetected with anti-*Ath*PEPCK1 antiserum (lower panel). B) Proteolysis of *Ath*PEPCK1 by recombinant *Ath*MC9. Reactions were done as described in the Materials and methods section, and aliquots were taken at the specified time-intervals. Then, samples were resolved in a 10% SDS-PAGE (upper panel), transferred to a nitrocellulose membrane and immunodetected with anti-*Ath*PEPCK1 antiserum (lower panel). C) Scheme of the *Ath*PEPCK1 truncated mutants constructed according to *Ath*MC9 recognition sites identified by Tisiatsiani *et al*. (2013).

To analyze the cleavage effect on *Ath*PEPCK1 activity, we cloned the gene coding for *Ath*MC9 from Arabidopsis seedlings and expressed the recombinant protein in *E. coli* cells. Recombinant *Ath*MC9 is expressed as a 37.1 kDa zymogen, which is auto-proteolyzed to produce the p10 (15.4 kDa) and p20 (21.73 kDa) subunits that conform the active *Ath*MC9 (Fig. S4; Vercammen *et al*., 2004). Incubation of *Ath*PEPCK1 with *Ath*MC9 for 1 h completely proteolyzed the former (Fig. 2B). The bands observed in Fig. 2B correspond to the theoretical fragments predicted according to the *Ath*MC9 recognition sites (K19 and R101; Fig. 2C) (Tsiatsiani *et al*., 2013). The activity of proteolyzed *Ath*PEPCK1 was slightly higher than the full-length protein, although the change was not statistically significant (data not shown).

To further study the effects of *Ath*MC9 cleavage on *Ath*PEPCK1 kinetics, we constructed two N-terminal truncated forms of *Ath*PEPCK1, Δ19 and Δ101. We expressed, purified, and characterized these mutants (Table 1, Figs S5-S6). In the carboxylation reaction, the *k*_cat_ of the Δ19 and Δ101 truncated enzymes were 1.5- and 1.7-fold higher than the WT, respectively; conversely, all enzymes had similar *k*_cat_ values in the reaction of decarboxylation (Table 1). Based on *K*_M_ values, both mutants have higher apparent affinities for nucleotides (ATP and ADP) and OAA than for PEP, like the WT enzyme (Table 1). The Δ19 mutant showed 2- and 6-fold higher apparent affinity for PEP and OAA than the WT, respectively. Contrarily, the Δ101 mutant showed 2- and 1.5-fold lower apparent affinity for PEP and OAA than the WT, respectively. Size exclusion chromatography revealed that both truncated forms are hexamers (Fig. S7), like the WT enzyme (Rojas *et al*., 2019).

**Table 1.**
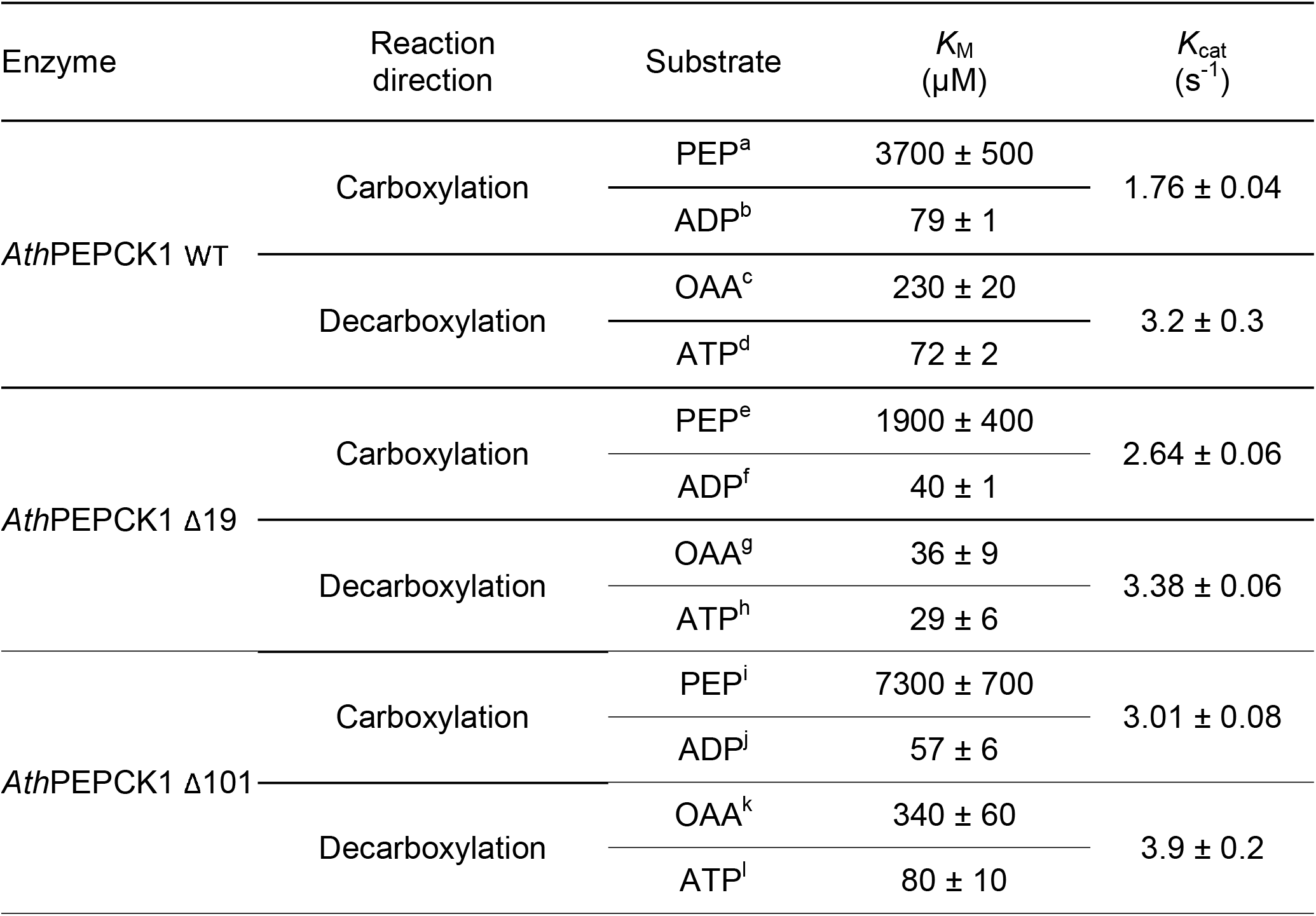
Kinetic parameters for *Ath*PEPCK1 and truncated mutants. Reactions in both directions of the reaction were performed as described in the Materials and methods section. Kinetic constants were calculated by fitting experimental data (Figs S5-S6) to the Michaelis-Menten equation. Reported values were provided by the fitting software. Fixed substrate concentrations were as follows: ^a^0.25 mM ADP, ^b^15 mM PEP, ^c^0.75 mM ATP, ^d^0.75 mM OAA, ^e^0.5 mM ADP, ^f^5 mM PEP, ^g^1.0 mM ATP, ^h^0.5 mM OAA, 0.5 mM ADP, ^j^15 mM PEP, ^k^0.5 mM ATP, and ^i^0.5 mM OAA. The parameters of the WT enzyme were taken from Rojas *et al*. (2019).

Plant PEPCKs are allosterically regulated by metabolites (Hatch and Mau, 1977; Leegood and Ap Rees, 1978; Burnell, 1986; Martín *et al*., 2011; Rojas *et al*., 2019). Interestingly, the response of the Δ19 and Δ101 mutants to different metabolites was altered compared with the full-length form. Both truncated mutants were more sensitive to PPi than the WT (Fig. 3). Sensitivity to shikimate was slightly increased in the Δ19 mutant compared with the WT, while the Δ101 mutant was almost insensitive to this metabolite (Fig. 3). Glc6P inhibited the WT enzyme and the Δ19 truncated form but, surprisingly, the Δ101 truncated form was 2-fold activated by the same metabolite (Fig. 3). Malate activated by 2- and 1.5-fold the WT enzyme and the Δ19 truncated form, respectively, while the Δ101 truncated form was insensitive to this metabolite (Fig. 3). To test whether malate still binds to the Δ101 mutant, we performed thermal-shift assays (Rojas *et al*., 2019). We found that malate produced a similar shift in the melting temperature (T_m_) of the WT, Δ19, and Δ101 truncated forms (Figs 4 and S8), suggesting this metabolite binds to all enzyme forms.

**Fig. 3.**
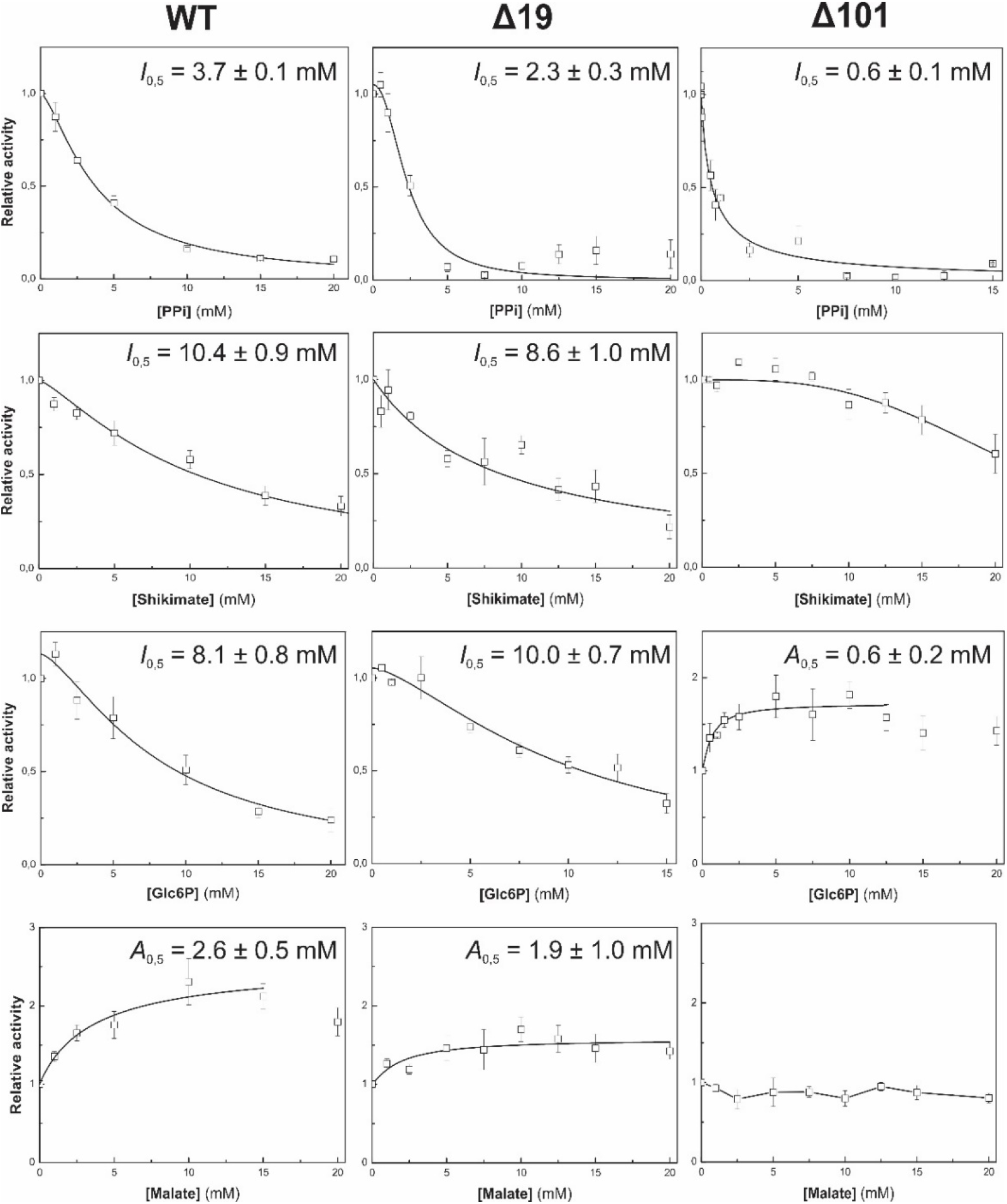
Analysis of allosteric effectors on *Ath*PEPCK1 and truncated mutants. Activity of *Ath*PEPCK1 WT (left panel), Δ19 (middle panel) and Δ101 (right panel) was measured in the direction of decarboxylation, as described in the Materials and methods section, using saturating substrate concentrations (0.75 mM OAA and 0.75 mM ATP for the WT, and 0.5 mM OAA and 0.3 mM ATP for the Δ19 and Δ101 mutants). In the case of malate, activity was measured in the carboxylation direction using saturating substrate concentrations (10 mM PEP and 0.13 mM ADP).

**Fig. 4.**
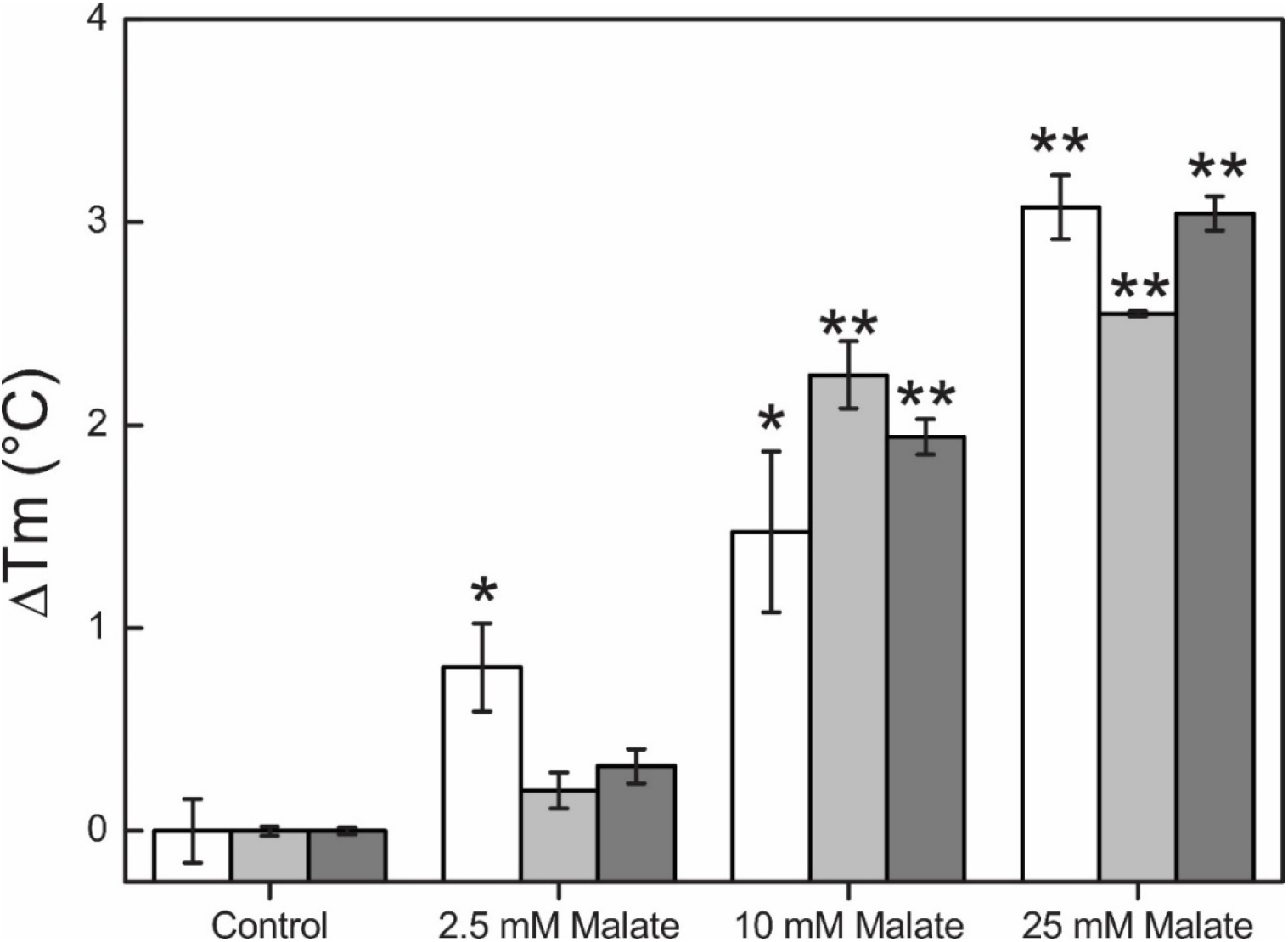
Thermal shift assays of *Ath*PEPCK1 and truncated mutants. Experiments were performed with the WT (white), Δ19 (light grey), and Δ101 (dark grey) enzymes in the presence of malate at the indicated concentrations. The shift in the melting temperature (T_m_) was calculated as described in the Materials and methods section. Data are the mean ± standard error of three technical replicates. * indicates a P-value < 0.05 and ** indicates a P-value < 0.01 using a t-test for two independent samples with a confidence level of 95%.

## Discussion

The regulation of enzymes by proteolysis is an emergent issue in plant biochemistry. The Arabidopsis genome codes for over 800 proteases with distinct temporal and tissue expression profiles (van der Hoorn, 2008; Tsiatsiani *et al*., 2012) but our knowledge on plant proteolytic cascades is still fragmentary (Paulus and Van der Hoorn, 2019). Understanding the regulation of the gluconeogenic pathway during seed germination is of critical importance, as seedling establishment has a direct impact on plant fitness and productivity (Graham, 2008). Based on this, we focused our studies on the proteolytic regulation of *Ath*PEPCK1, the pacemaker enzyme of plant gluconeogenesis (Penfield *et al*., 2004, 2012; Eastmond *et al*., 2015).

Some authors have observed that plant enzymes usually contain N- and C-terminal extensions compared to their bacterial or cyanobacterial counterparts. Such extensions are generally susceptible to post-translational modifications (Lepiniec *et al*., 1993; Ocheretina *et al*., 1993; Walker and Leegood, 1995; Furumoto *et al*., 1999). These extensions might represent regulation modules acquired during evolution to accomplish complex regulations, as plants must respond to ever-changing environmental conditions. The first observation of a discrete proteolytic cleavage at the N-terminus of a plant PEPCK was reported by Walker *et al*. (1995). These authors demonstrated that PEPCK was proteolyzed in crude extracts obtained at neutral pH, which could be prevented by making extractions at alkaline pH (Walker *et al*., 1995). We found similar results in Arabidopsis, as native extractions at alkaline pH prevented further PEPCK degradation (Figs 2A and S3). Cleavage of the cucumber PEPCK did not significantly alter its activity (Walker and Leegood, 1995). Therefore, the authors hypothesized that the N-terminal extension would confer unique regulatory properties to the enzyme, not located in the smaller bacterial versions. This model was supported by the fact that plant PEPCK is phosphorylated at the N-terminus, which in turn inhibits the activity of the enzyme (Walker and Leegood, 1995; Leegood and Walker, 1996, 2003; Walker *et al*., 1997, 2002; Bailey *et al*., 2007; Chao *et al*., 2014; Shen *et al*., 2017).

In our studies with Arabidopsis seedlings, we found that *Ath*PEPCK1 is subject to proteolysis around 24-48 HAI, when the level of protein reaches a maximum (Figs 1A-C). We measured PEPCK carboxylation activity in these samples, but we did not find the peak of the activity described by Malone *et al*. (2007); instead, we observed a gradual increase of PEPCK activity (~2-fold), coincident with the peak of protein accumulation (Fig. 1C). These differences might be originated by distinct extraction conditions, which might lead to different enzyme populations. By making denaturing protein extractions, we found that the shorter versions are present in Arabidopsis germinating seedlings, as reported for PEPCK from pineapple leaves (Martín *et al*., 2011).

A large-scale study conducted by Tsiatsiani *et al*. (2013) identified *Ath*PEPCK1 as a target of *Ath*MC9. Crude extracts of *mc9* knock-out mutants and MC9 overexpressing lines showed decreased and increased PEPCK carboxylase activity, respectively (Tsiatsiani *et al*., 2013). The recombinant *Ath*PEPCK1 truncated mutants (Δ19 and Δ101) showed similar decarboxylation activity than the WT enzyme. In comparison, the carboxylation reaction was slightly increased in both truncated forms compared to the WT enzyme (Table 1). The *K*_M_ values for the substrates of the truncated mutants were in the same range as those determined for the short version of the pineapple PEPCK (Martín *et al*., 2011). The *K*_M_ for PEP of the Δ19 mutant was 2-fold lower than the WT enzyme; similarly, the short version of pineapple PEPCK has a 10-fold lower *K*_M_ for PEP than the entire isoform (Daley *et al*., 1977; Martín *et al*., 2011).

A key characteristic of the *Ath*PEPCK1 truncated mutants is that allosteric regulation by metabolites differs from that observed for the WT enzyme. Particularly, Glc6P is an inhibitor of the WT enzyme, but a soft activator of the Δ101 truncated form; whereas malate activates the WT enzyme and has no effect on the Δ101 truncated form (Fig. 3). These characteristics reinforce the idea that the N-terminal extension confers regulatory properties to plant PEPCK. These results agree with the findings of Furumoto *et al*. (1999), who treated maize PEPCK with enterokinase under controlled conditions to cleave the N-terminus. The proteolyzed enzyme showed 2-fold higher activity and was inhibited by 3PGA, while the full-length protein was only slightly affected by this metabolite (Furumoto *et al*., 1999). In line with our findings, Martín *et al*. (Martín *et al*., 2011) showed that the small version of pineapple PEPCK is not affected by Glc6P or L-malate in the decarboxylation direction of the reaction; unfortunately, we do not know if these metabolites have any effect on the full-length form of the pineapple enzyme, as it could not be purified and characterized (Martín *et al*., 2011).

The proteolytic regulation of *Ath*PEPCK1 might be part of a mechanism to regulate its levels and/or activity during the sink-to-source transition. During germination, when carbon is obtained from lipids and amino acids, the levels of malate increase, thus activating *Ath*PEPCK1 and the flux of carbon into gluconeogenesis (Fig. 5). Once the photosynthetic apparatus is developed, gluconeogenesis is replaced by glycolysis. At this stage, reduced carbon is obtained from the Benson-Calvin-Basham cycle, the levels of hexose-phosphates increase and PEPCK activity diminishes approximately 10-fold, from ~0.5 U g FW^−1^ in germinating seedlings to ~0.05 U g FW^−1^ in mature leaves (Fig. 1C and Malone *et al*., 2007). At this point, *Ath*MC9 might generate the shorter *Ath*PEPCK1 isoforms needed for the new metabolic scenario; i.e., to provide intermediates to the tricarboxylic acid cycle (Fig. 5).

**Fig. 5.**
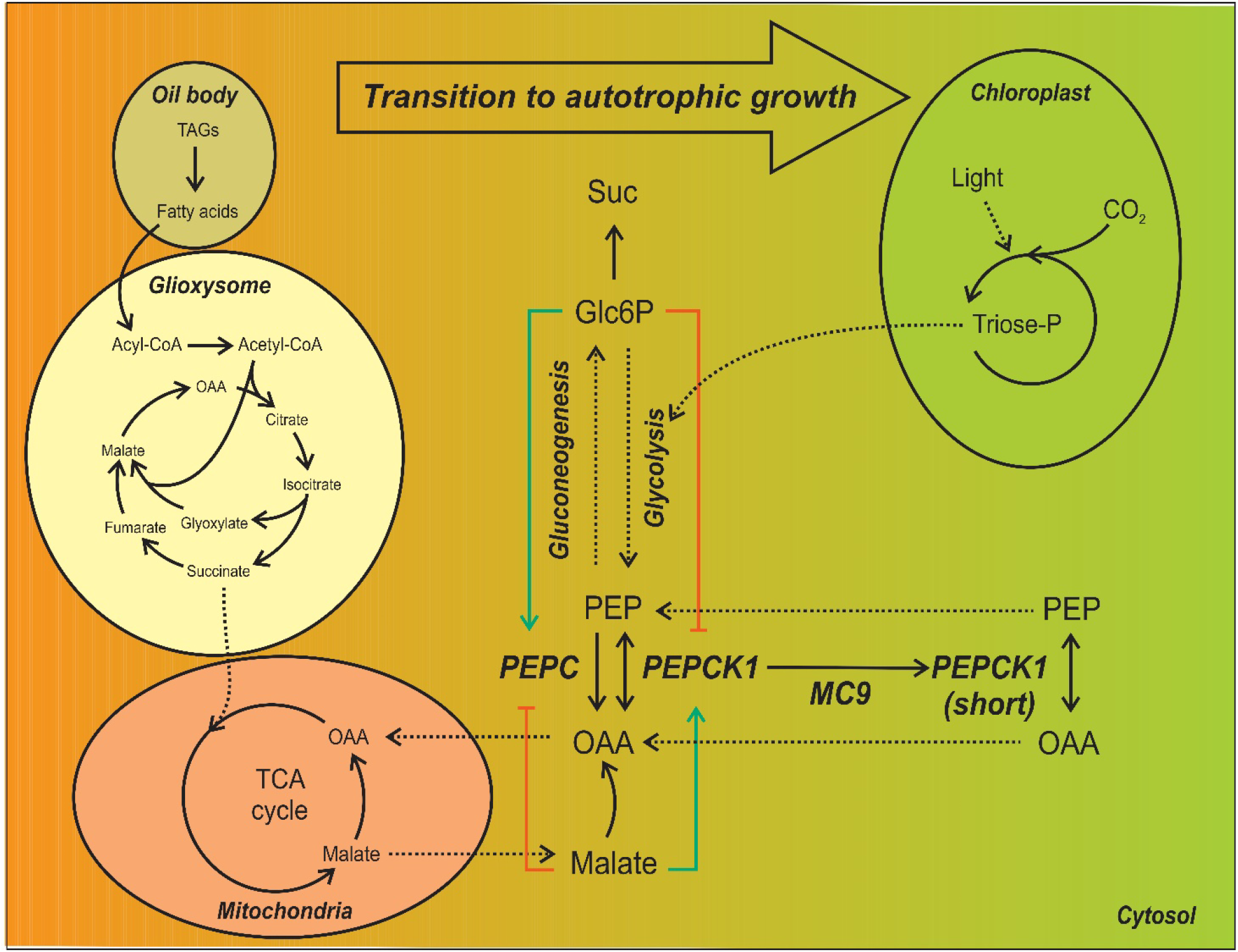
Regulation of *Ath*PEPCK1 during the sink-to-source transition. During the transition of the seedling from heterotrophic (orange) to autotrophic (green) growth, *Ath*PEPCK1 is cleaved at the N-terminus by *Ath*MC9, leading to shorter enzyme forms, which are catalytic but insensitive to regulation by malate and Glc6P. Blue lines, activation; red lines, inhibition.

## Abbreviations

*Ath*MC9: *Arabidopsis thaliana* metacaspase-9
*Ath*PEPCK1: *Arabidopsis thaliana* phospho*enol*pyruvate carboxykinase-1
Glc6P: glucose 6-phosphate
OAA: oxaloacetic acid
PEP: phospho*enol*pyruvate
PPi: inorganic pyrophosphate

## Acknowledgements

This work was supported by grants from ANPCyT (PICT-2017-1515 and PICT-2018-00929 to AAI, and PICT-2018-00865 to CMF) and UNL (CAI+D). CMF is funded by the Max Planck Society (Partner Group for Plant Biochemistry). BER is a Fellow from CONICET, CMF and AAI are Researchers from the same Institution.

